# Integrated analysis of oral tongue squamous cell carcinoma identifies key variants and pathways linked to risk habits, HPV, clinical parameters and tumor recurrence

**DOI:** 10.1101/028845

**Authors:** Neeraja M Krishnan, Saurabh Gupta, Vinayak Palve, Linu Varghese, Swetansu Pattnaik, Prachi Jain, Costerwell Khyriem, Arun K Hariharan, Kunal Dhas, Jayalakshmi Nair, Manisha Pareek, Venkatesh K Prasad, Gangotri Siddappa, Amritha Suresh, Vikram D Kekatpure, Moni Abraham Kuriakose, Binay Panda

**Affiliations:** Ganit Labs, Bio-IT Centre, Institute of Bioinformatics and Applied Biotechnology, Bangalore, India; Integrated Head and Neck Oncology Program, Mazumdar Shaw Centre for Translational Research, Bangalore, India; Head and Neck Oncology, Mazumdar Shaw Medical Centre, Bangalore, India; Strand Life Sciences, Bangalore, India

**Keywords:** oral tongue squamous cell carcinoma, somatic variants, gene expression, CNV, LOH, tumor recurrence, *CASP8*, *TP53*, *RASA1*, HPV

## Abstract

Oral tongue squamous cell carcinomas (OTSCC) are a homogenous group of tumors characterized by aggressive behavior, early spread to lymph nodes and a higher rate of regional failure. Additionally, the incidence of OTSCC among younger population (<50yrs) is on a rise; many of who lack the typical associated risk factors of alcohol and/or tobacco exposure. We present data on SNVs, indels, regions with LOH, and CNVs from fifty-paired oral tongue primary tumors and link the significant somatic variants with clinical parameters, epidemiological factors including HPV infection and tumor recurrence. Apart from the frequent somatic variants harbored in *TP53, CASP8, RASA1, NOTCH* and *CDKN2A* genes, significant amplifications and/or deletions were detected in chromosomes 6-9, and 11 in the tumors. Variants in *CASP8* and *CDKN2A* were mutually exclusive. *CDKN2A, P1K3CA, RASA1* and *DMD* variants were exclusively linked to smoking, chewing, HPV infection and tumor stage. We also performed whole-genome gene expression study that identified matrix metalloproteases to be highly expressed in tumors and linked pathways involving arachidonic acid and NF-κ-B to habits and distant metastasis, respectively. Functional knockdown studies in cell lines demonstrated the role of *CASP8* in HPV-negative OTSCC cell line. Finally, we identified a 38-gene minimal signature that predicts tumor recurrence using an ensemble machine learning method. Taken together, this study links molecular signatures to various clinical and epidemiological factors in a homogeneous tumor population with a relatively high HPV prevalence.

## Introduction

Squamous cell carcinomas of head and neck (HNSCC) are the sixth leading cause of cancer worldwide [1]. Tumors of head and neck region are heterogeneous in nature with different incidences, mortalities and prognosis for different subsites and accounts for almost 30% of all cancer cases in India [2]. Oral cancer is the most common subtype of head and neck cancers in humans, with a worldwide incidence in >300,000 cases. The disease is an important cause of death and morbidity, with a 5-year survival of less than 50% [1, 2]. Recent studies have identified various genetic changes in many subsites of head and neck using high-throughput sequencing assays and computational methods [3-7]. Such multi-tiered approaches using the exomes, genomes, transcriptomes and methylomes from different squamous cell carcinomas have generated data on key variants and in some cases, their biological significance, aiding our understanding of disease progression. Some of the above sequencing studies have identified key somatic variants and linked them with patient stratification and prognostication. This, along with the associated epidemiology, enables one to look beyond the discovery of driver mutations, and identify predictive signatures in HNSCC.

A previous study from the cancer genome atlas (TCGA) consortium with HNSCC patients (N = 279) identified somatic mutations in *TP53, CDKN2A, FAT1, P1K3CA, NOTCH1, KMT2D* and *NSD1* at a frequency greater than 10% [7]. Additionally, the TCGA study identified loss of *TRAF3* gene, amplification of *E2F1* in human papilloma virus (HPV)-positive oropharyngeal tumors, along with mutations in *P1K3CA, CASP8* and *HRAS,* and co-amplifications of the regions 11q13 (harboring *CCND1, FADD* and *CTTN*) and 11q22 (harboring *B1RC2* and *YAP1*), in HPV-negative tumors, described to play an important role in pathogenesis and tumor development [7]. Chromosomal losses at 3p and 8p, and gains at 3q, 5p and 8q were also observed in HNSCC [7]. Tumors originating in the anterior/oral part of tongue or, oral tongue squamous cell carcinoma (OTSCC) tend to be different from those at other subsites as oral tongue tumors are associated more with younger patients [8] and spread early to lymph nodes [9]. Additionally oral tongue tumors have a higher regional failure compared to gingivo-buccal cases [10] in oral cavity. Tobacco (both chewing and smoking) and alcohol are common risk factors for this group of tumors among older patients [8]. The role of HPV, both as an etiological agent and/or risk factor along with its role as a good prognostic marker in OTSCC, unlike in oropharyngeal tumors, is currently uncertain. It remains to be explored whether HPV acts as an etiological agent in the development of oral tongue tumors or simply represents a super infection in patients. Additionally, HPV infection status currently does not influence disease management in OTSCC.

Here, we present data towards a comprehensive molecular characterization of OTSCC. We performed exome sequencing, whole-genome gene expression, and genotyping arrays using fifty primary tumors along with their matched control samples, towards identification of somatic variants (mutations and indels), significantly up- and down-regulated genes, loss of heterozygosity (LOH) and copy number variations (CNVs). We integrated all the molecular data along with the clinical parameters and epidemiology such as tumor stage, nodal status, HPV infection, risk habits and tumor recurrence to interpret the effect of changes in the process of cancer development in oral tongue. We identified significant somatic variations in *TP53* (38%), *RASA1* (8%), *CASP8* (8%), *CDKN2A* (6%), *NOTCH1* (4%), *NOTCH2* (4%), and *P1K3CA* (4%) from the exome sequencing study in OTSCC. The key variants were validated using an additional set of primary tumor samples. Variants in *TP53* and *NOTCH1* were found in mutually exclusive sets of tumors. Additionally, we found frequent aberrations in chromosomes 6-9, and 11 in tumor samples. We observed a strong association between somatic variations in some key genes with one or more risk habits; for example, *CDKN2A* and *P1K3CA* with smoking; *CASP8* with consuming alcohol and chewing tobacco; *RASA1* with chewing and tumor stage, and HPV infection, along with *DMD* and *P1K3CA.* From the gene expression analysis, we found matrix metalloproteases (MMPs) to be highly expressed in OTSCC. Pathway analysis identified Procaspase-8, Notch, Wnt, p53, extracellular matrix (ECM)-receptor interaction, JAK-STAT and PPAR to be some of the significantly altered pathways in OTSCC. We implemented an ensemble machine learning method [11] and identified a minimal gene signature set that distinguished a group of tumors with loco-regional recurrence from the non-recurrent set. Finally, we performed functional analysis of *CASP8* gene in HPV-negative and HPV-positive OTSCC cell lines to establish its role in the process of tumor development.

## Results

### Habits, clinical parameters and epidemiology

We collected tumor and matched control (adjacent normal and/or lymphocytes) samples from 50 patients diagnosed with OTSCC, with informed consent. Data from patient habits, epidemiology and clinical parameters are presented in Figure 1A and Additional file 1A. About two-thirds of the patients (N = 31) included in our study were in the younger age group (≤50yrs), with 20% female patients in the total pool. Approximately, 70% of the patients were positive for at least one risk habit, namely, smoking, alcohol consumption or chewing tobacco (33% of patients smoked tobacco, 40% consumed alcohol and 42% chewed tobacco). HPV infection status in the primary tumors was established with type-specific qPCR or HPV16 digital PCR. Thirty-three percentage of the patients were deceased at the time of completing the analysis. About 60% of the tumors were moderately differentiated, 25% well differentiated and the rest were poorly differentiated. Among the patients recruited, 60% were node-positive, 70% had no recurrence, 9% had distant metastasis and 24% had loco-regional recurrence at the time of completing the analysis. The mean and median follow up duration for patients were nearly 30 months and 21 months respectively. About 27% of the tumors were early stage tumors (T1N0M0 and T2N0M0) and the rest 73% were late stage tumors (tumors belonging to the rest of the TNM stage).

**Figure 1.**
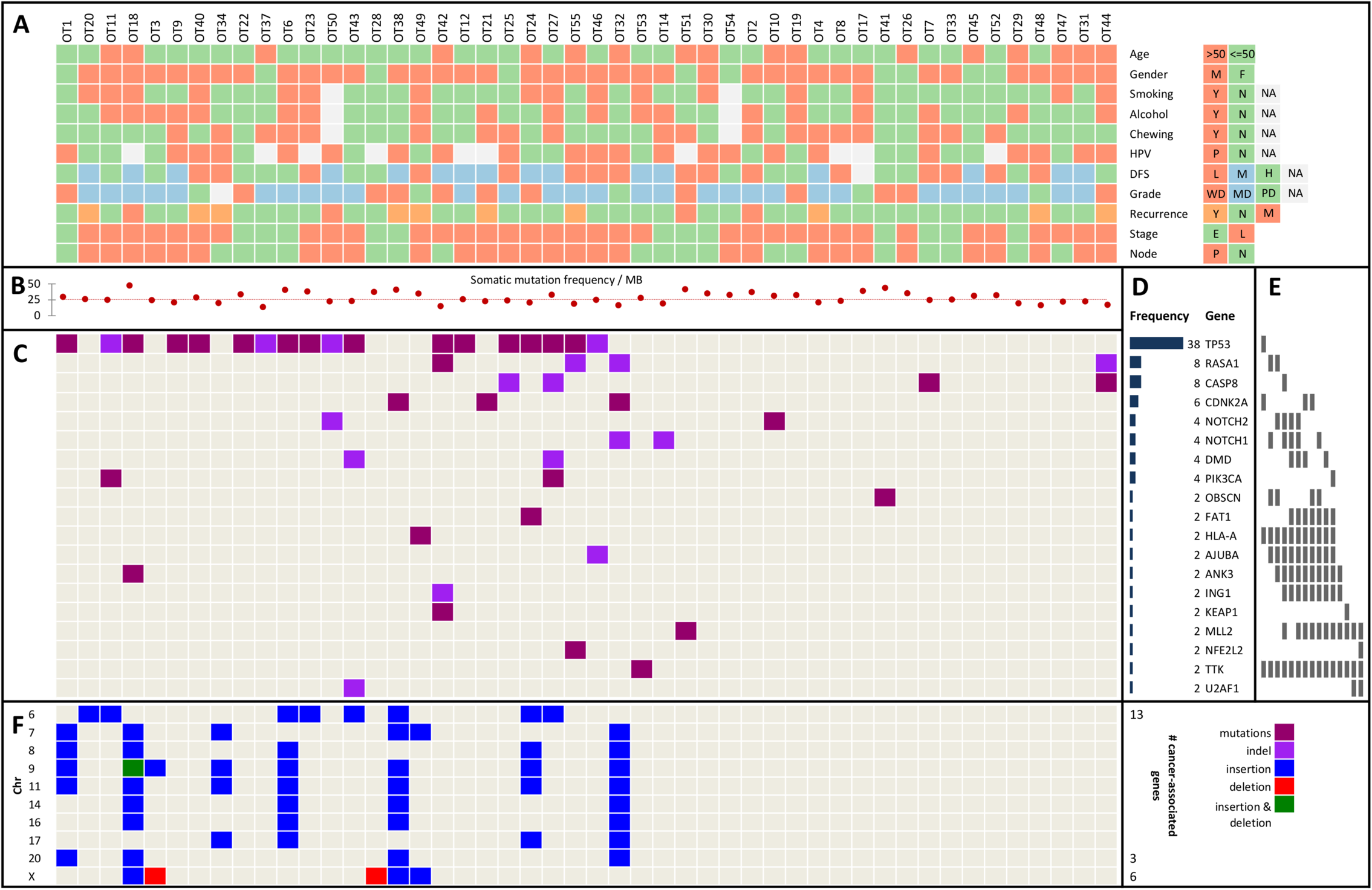
Key variants in OTSCC and their relationship with habits, clinical and epidemiological parameters. A. The OTSCC samples are represented in color-codes with their corresponding status on; node (P: positive, N: negative); stage (E: early, L: late), recurrence (Y: loco-regionally recurrent, N: non-recurrent and M: distant metastatic); grade (WD: well-differentiated, MD: moderately-differentiated and PD: poorly-differentiated); disease-free survival or DFS (L: low/≤12mo, M: mid/12-24mo and H: high/>24mo); HPV (P: positive and N: negative); and habits (chewing, alcohol and smoking, Y: yes and N: no). B. Somatic mutation frequency per megabase (MB) is represented as scatterplot with the median point as a fine dotted line. C. Genes with significant somatic variants. D. Frequency histogram of nineteen cancer-associated genes bearing somatic missense and nonsense variants (mutations and indels). E. Columns representing mutually exclusive sets of genes. F. Significant copy number insertions and deletions (CNVs), alongside the chromosome cytobands (the numbers of cancer-associated genes within each cytoband are listed on the right).

### Discovery and validation of significant somatic variants and their relationship with other parameters

We re-discovered variants, as described previously [12] using whole-genome arrays, to validate the variant call accuracy as obtained from the exome sequencing data. We validated ~99% of the SNPs discovered from Illumina sequencing in both the tumor and matched control samples (Additional file 2). After filtering and annotation, we identified 19 cancer-associated genes bearing significantly altered somatic variants in OTSCC (Figure 1D). These were validated using Sanger sequencing in two sets of samples, one using the same tumor-control pairs used in the exome sequencing (the discovery set, Additional file 1A) and second, using an additional 36-60 primary tumors (validation set, Additional file 1B) for genes altered in ≥ 5% of the tumor samples. All the *TP53* variants were validated in the discovery set. Three out of the four variants were validated for *CASP8.* The mutant alleles for the heterozygous variants in *HLA-A, OBSCN, ING1, TTK* and *U2AF1* discovered by exome sequencing were difficult to interpret from the results of the validation using Sanger sequencing as they were present at a very low frequency (Additional file 3). Combining data from the validation set; the mutation frequencies for *RASA1* and *CDKN2A* rose significantly to 10.71% and 16.47% in primary tumors respectively but those for *TP53* and *CASP8* remained largely unchanged (Additional file 3).

The somatic mutation frequency per MB ranged from 10-45 with a median around 25 (Figure 1B). The median value for transition to transversion (ti/tv) ratio for both the tumor and its matched control samples was ~2.5 (Additional file 4). Overall, *T->C* changes were most frequent, followed by *G->A* and then *T->G.* Habits (smoking and alcohol consumption), nodal status, HPV infection, tumor grade and stage had no significant impact on the distribution of these nucleotides (Additional file 5). We used the workflow described in the *Methods* section to identify somatic mutations and indels in tumor samples following which we used three functional tools, IntOGen [19], MutSigCV [21] and MuSiC2 [22] for variant interpretations (Additional file 6). In order to identify genes harboring significant variants, we used the intersection of these tools, following the criteria that the somatic variants be callable in the matched control sample and present in a single sequencing read in the control sample. This resulted in a final list of 19 cancer-associated genes (Figure 1C), which were divided into three categories with varying mutation frequencies (Figure 1D). The three frequency tiers were ≥ 30% *(TP53),* 6-30% *(RASA1, CASP8* and *CDKN2A)* and 2-5% *(NOTCH1, NOTCH2, DMD* and *P1K3CA* were prominent among them).

Next, we looked for mutual exclusivity of finding somatic variants in the genes and found that many of these genes harbor variants in a mutually exclusive manner across samples (Figure 1E), suggesting the possibility that there might be some common pathway(s) involved in the development of OTSCC. We observed mutual exclusivity among somatic variants in *NOTCH1* and *NOTCH2* genes, and expanded this finding to identifying 15 such mutually exclusive sets (Figure 1E). Among them, *CDKN2A, HLA-A* and *TTK* form a mutually exclusive set with *TP53; RASA1, OBSCN, HLA-A, AJUBA* and *TTK* are mutually exclusive with either *NOTCH1* alone, or *NOTCH2* and *ANK3* together; *NOTCH1, NOTCH2, HLA-A, AJUBA, ANK3, TTK, MLL2, ING1* or *KEAP1,* are mutually exclusive with *CASP8* alone, or *FAT1* and *DMD* together; *FAT1, HLA-A, AJUBA, ANK3, TTK*, *MLL2,1NG1* or *KEAP1,* are mutually exclusive with *P1K3CA* or *DMD* or *NOTCH1* and *OBSCN,* or *CDKN2A* and *OBSCN*; *U2AF1, MLL2* and *TTK* form a small mutually exclusive set. We juxtaposed the positions of the somatic variants from final list of all 19 genes (Additional file 7) detected in OTSCC against those found in the TCGA data using the cBioPortal. We found that the somatic variants in OTSCC were in the same domains where mutations were observed earlier in many of the genes (Additional file 7).

Copy Number Variation (CNV) analyses using data from the whole-genome SNP genotyping arrays revealed a large chunk of chromosome 9, bearing cancer-associated genes like *CDKN2A, NF1* and *MRPL4,* to be affected in about 17% of the tumors (Figure 1F and Additional file 8). We found several CNVs of short stretches (in low kb range) within chromosomes 6-8, 11, 17 and X in many tumors.

### Linking habits, HPV infection, nodal status, tumor grade and recurrence, with genes harboring somatic variants and the associated pathways

We further classified the 19 cancer-associated genes from the previous analyses and linked those with habits, clinical parameters and HPV infection. Among the genes harboring significant somatic variants, we found *CDKN2A* to be mutated only in the never smokers and past smokers, *PIK3CA* to be mutated only in the smokers, and *TP53* to be mutated at a 20% greater frequency in the smokers, *CASP8* has a 12% greater frequency in those that consumed alcohol or chewed tobacco. *RASA1* was exclusively mutated only in the non-chewers. (Figure 2A). HPV-negative patients harbored somatic variants in *DMD* and *PIK3CA,* while HPV-positive patients alone had somatic variants in *RASA1.* Only the moderate-and well-differentiated tumor samples harbored variants in *CASP8*, while *NOTCH1* was mutated largely in the poorly-differentiated tumors. Node-positive tumors had a 19% greater occurrence of *TP53* variants. Somatic variants in *RASA1* occurred exclusively in the late stage tumors (Figure 2A). We further studied the association of affected cancer-related signaling pathways with habits and clinical parameters, and found that recurrence and HPV infection had the highest impact (Figure 2B). The Procaspase-8 activation, Notch, p53 and Wnt signaling pathways were linked most with many of the clinical parameters, HPV infection and habits (Figure 2B).

**Figure 2.**
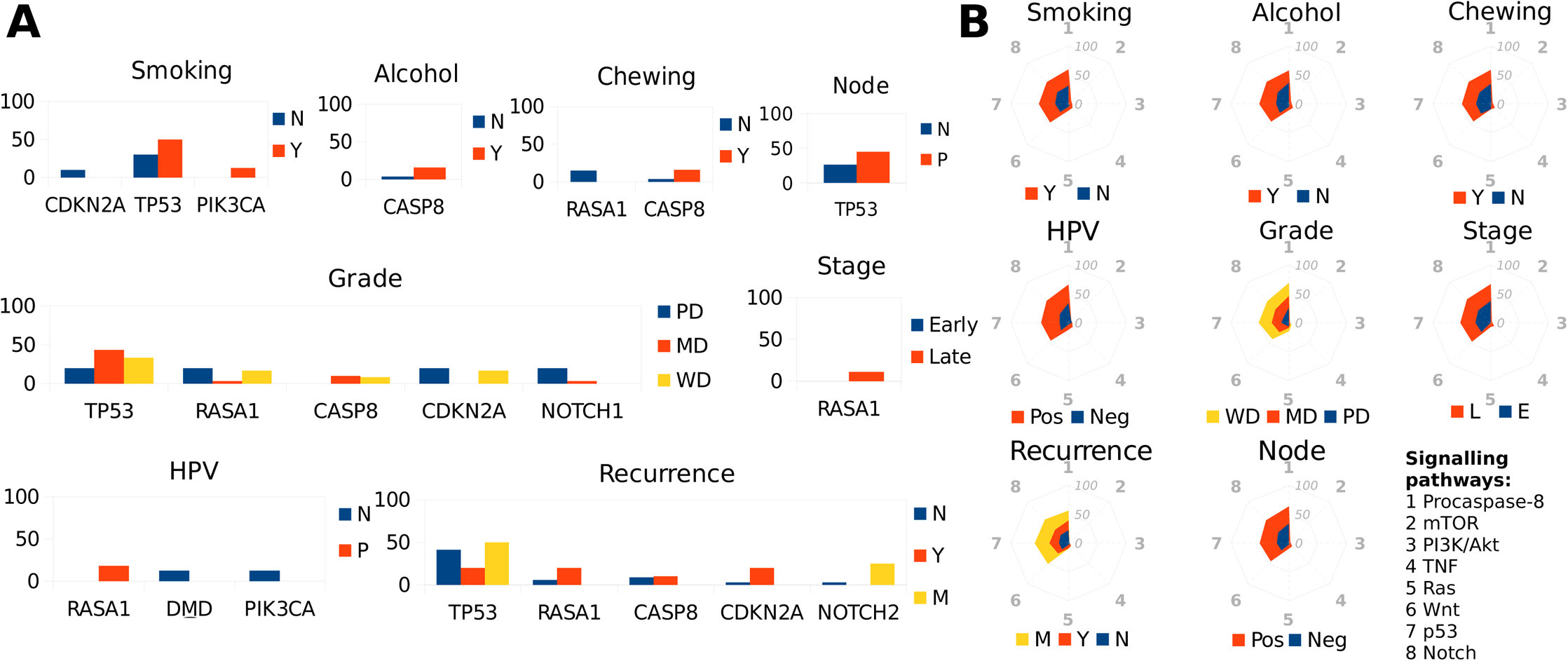
Relationship between genes harboring somatic variants with clinical-, epidemiological parameters and signaling pathways. A. Histograms showing relationship between genes with significant somatic variants and various clinical and epidemiological parameters,.for genes solely mutated in one of the clinical or epidemiological categories, or those mutated at a > = 5% frequency between two categories. B. Stack net charts of relative patient fraction (%) for each of the eight cancer-associated signaling pathways and their relationship with various clinical and epidemiological parameters.

### Differentially expressed genes in OTSCC

Significant (q val ≤ 0.05) differentially expressed genes with a fold change of at least 1.5 revealed a consistent pattern of differential expression across the tumor samples (21 up-and 23 down-regulated genes, Figure 3A and Additional file 9). Genes in PPAR signaling- (e.g., *MMP1*) and ECM-receptor interaction pathways *(LAMC2* and *SPP1)* were up-regulated and *CRNN, APOD, SCARA5* and *RERGL* were down-regulated in a majority of tumors (Figure 3A). Next, we studied the pathways involving genes with aberrant expression and their link with HPV infection and other clinical parameters. Genes in the arachidonic acid metabolism and Toll-like receptors were differentially expressed in patients with no smoking history (never smokers or past smokers) and alcohol habits (Figure 3B). *SERP1NE1* (a gene in HIF-1 signaling pathway) was differentially expressed in patients that are habits-negative. The NF-κ-B signaling pathway was differentially expressed only in metastasized tumors.

**Figure 3.**
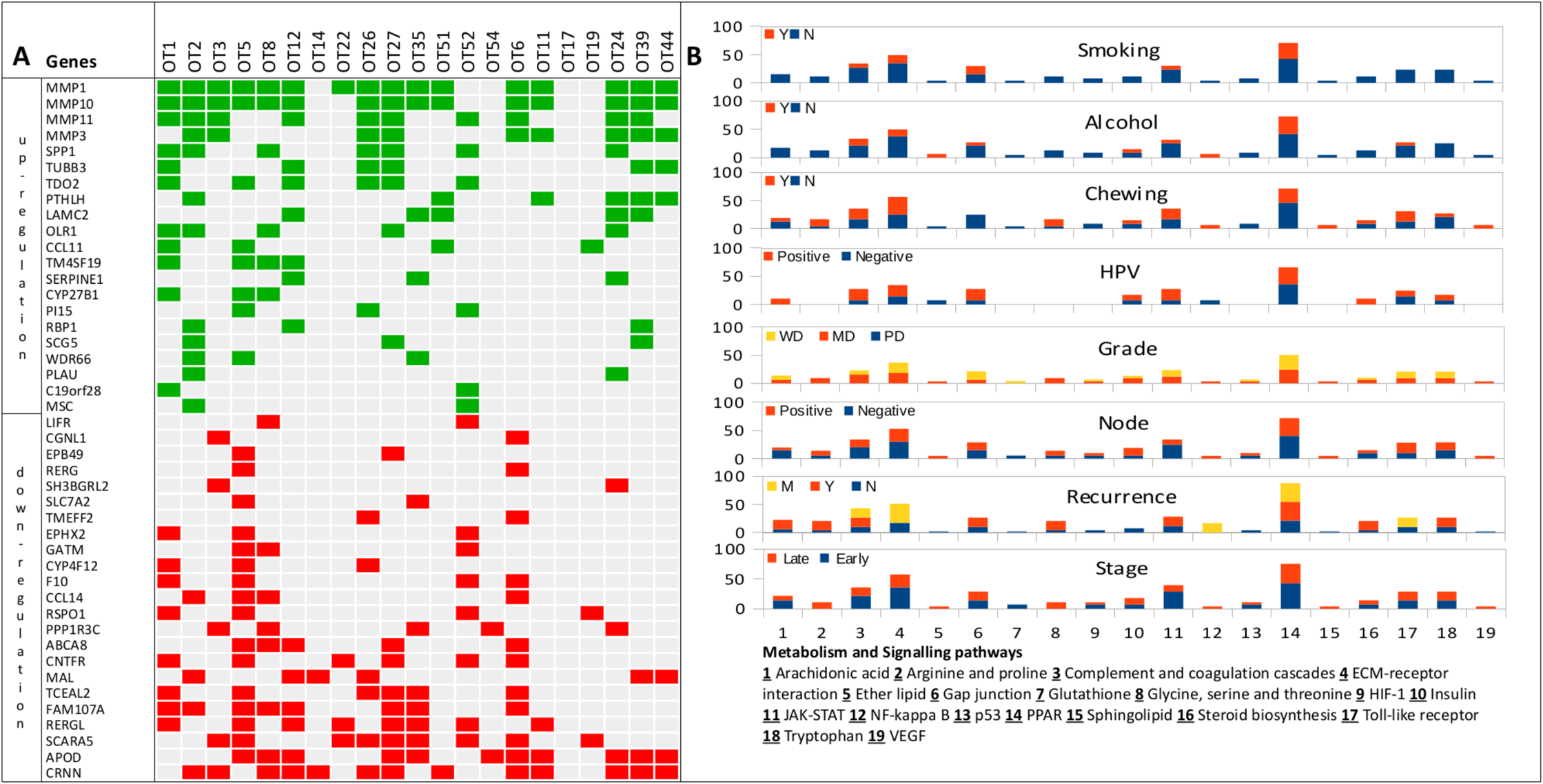
Differentially expressed genes, affected pathways and their relationship with clinical and epidemiological parameters. A. Expression changes (green – up- and red – down-regulation) representing significantly differentially expressed genes in tumors. B. Stacked histograms representing relative patient fraction (%) for each of the 19 cancer-associated pathways and their relationship with clinical and epidemiological parameters.

### Functional studies with CASP8 in OTSCC cell lines

*CASP8* is mutated in a significant number of oral tongue tumors [this study, 5, 7]. Caspase-8 is an important and versatile protein that plays a role in both apoptotic (extrinsic or death receptor-mediated) and non-apoptotic processes [13, 14]. We studied the functional consequences of *CASP8* knockdown through a siRNA-mediated method in an HPV-positive UM:SCC-47 [15] and an HPV-negative UPCI:SCC040 [16] OTSCC cell lines. Prior to the functional assay, the concentration of siRNA required for silencing, extent of *CASP8* knockdown and cisplatin sensitivity (IC_50_) in both these cell lines was tested (Additional file 10). The invasion of cells was greater in both UM:SCC-47 and UPCI:SCC040 cell lines when *CASP8* was knocked down (Figure 4A). To analyze the effect of caspase-8 on the migration property of cells, scratches were made on the confluent monolayer of cells and the wound closure area was measured at different time points (0hr, 15hr, 23hr & 42hr, Figure 4B). The wound closure was faster in *CASP8* knockdown HPV-negative cells compared to the HPV-positive cells. At 15hr, 23hr and 48hrs, about 65%, 90% and 100% of the wound got closed respectively in the HPV-negative cell line compared to 50%, 70% and 85% respectively during the same time period in the HPV-positive cell lines (Figure 4B). siRNA knockdown of *CASP8* rescued the chemo-sensitivity caused by cisplatin treatment as evident by the MTT (3-(4,5-dimethylthiazol-2-yl)-2,5-diphenyltetrazolium bromide) survival assay (Figure 4C). Interestingly, we found that the extent of rescue is greater in the HPV-negative cell line compared to the HPV16-positive one.

**Figure 4:**
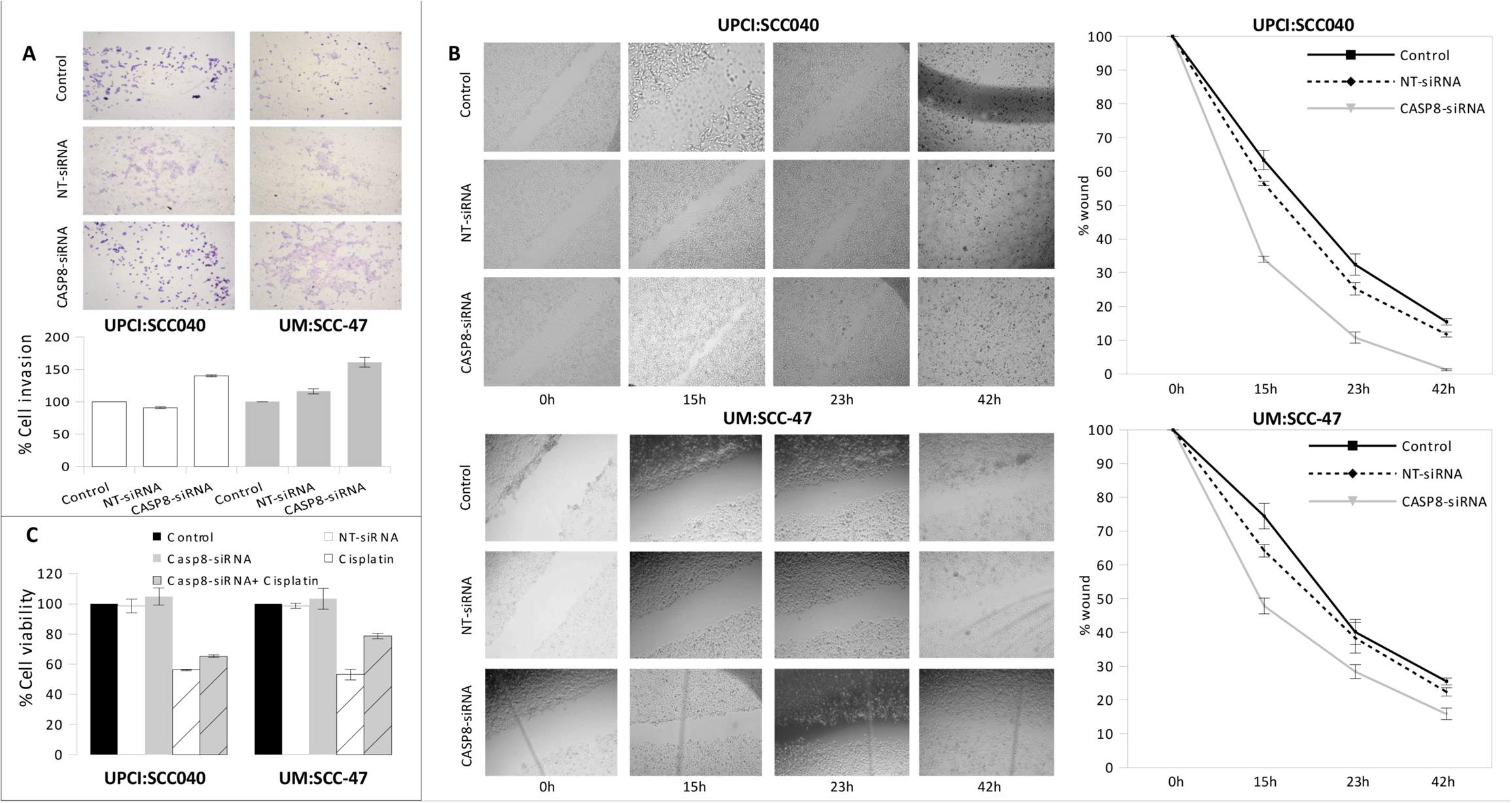
Role of *CASP8* in HPV-positive and HPV-negative OTSCC cell lines. Results from A. Matrigel cell invasion assay (plotted with respect to the control cells), B. Wound healing assay, and C. MTT cell survival assay (plotted with respect to the control cells) in UPCI:SCC040 (HPV-negative) and UM:SCC-47 (HPV-positive) cell lines.

### Tumor recurrence prediction using random forests

After cataloging the significantly altered genes in OTSCC, we wanted to see whether there is a relationship between the altered genes and loco-regional recurrence of tumors and metastasis. In order to do this, we used an ensemble machine learning method implemented by variable elimination using random forests [11] (Figure 5). We used multiple testing correction and the 0.632 bootstrapping method [17] to estimate false positives. We discovered a 38-gene minimal signature that discriminated between the non-recurring, loco-regionally recurring and distant metastatic tumors (Figure 5). The .632+ bootstrap errors, indicative of prediction specificity, varied across non-recurrent, recurrent and distant metastatic tumors. The median error was low (0.03) and intermediate (0.3) for the non-recurrent and the loco-regionally recurrent categories respectively but was relatively higher (1.0) for the metastatic tumors. The errors were proportional to the number of representative samples within each category.

**Figure 5.**
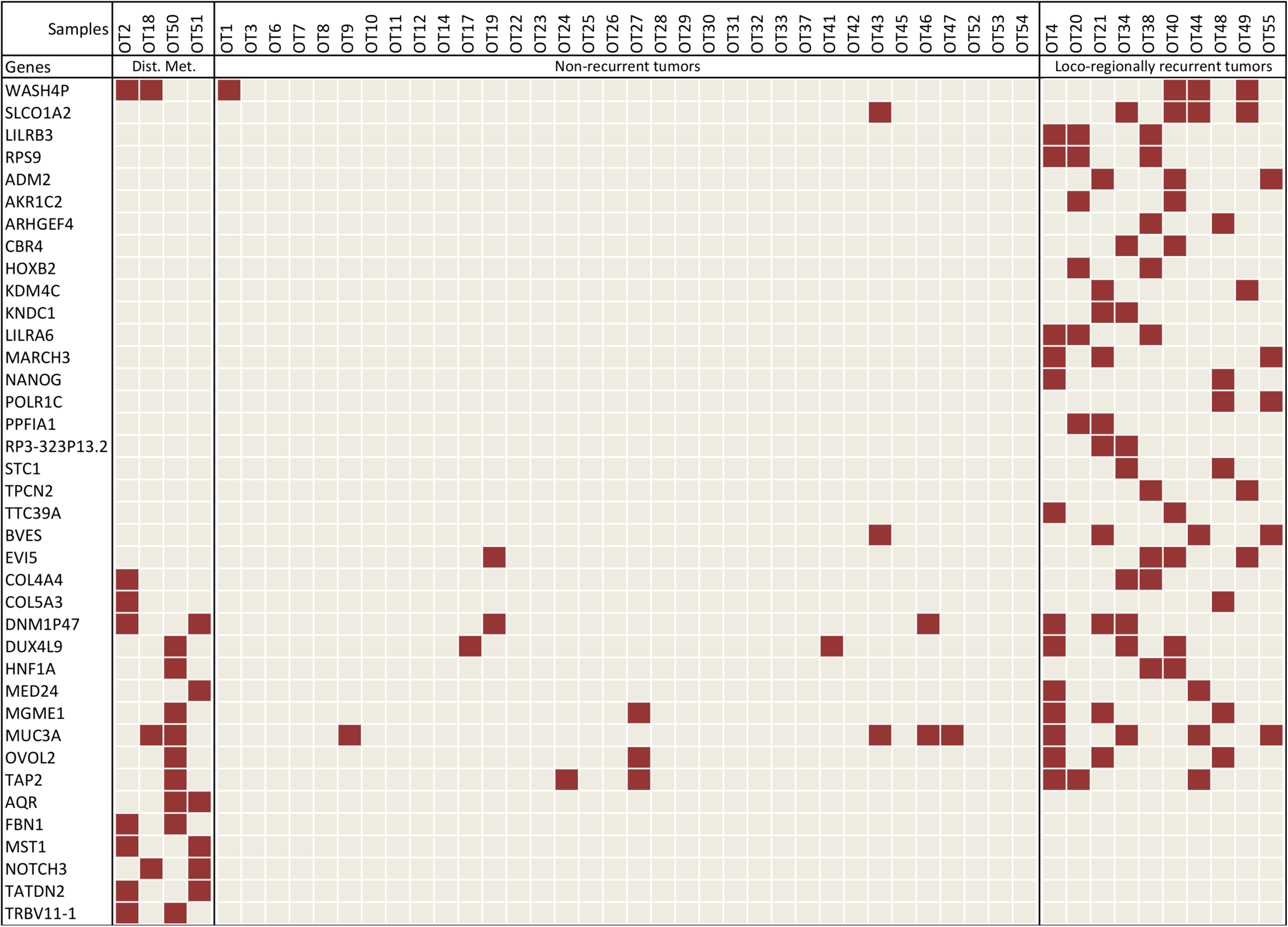
A minimal gene signature for tumor recurrence. Genes harboring somatic variants (in color) that are a part of the minimal signature set for tumor recurrence derived from random forest analyses are used.

### Major signaling pathways implicated in OTSCC

We looked at significant pathways altered in OTSCC, taking into account all the molecular changes in tumors and found apoptosis, HIF, Notch, mTOR, p53, PI3K/Akt, Wnt and Ras to be some of the key signaling pathways affected in OTSCC (Figure 6). In addition, histone methylation, cell cycle/immunity and mRNA splicing processes were also affected. The complete list of pathways is provided in Additional file 11.

**Figure 6.**
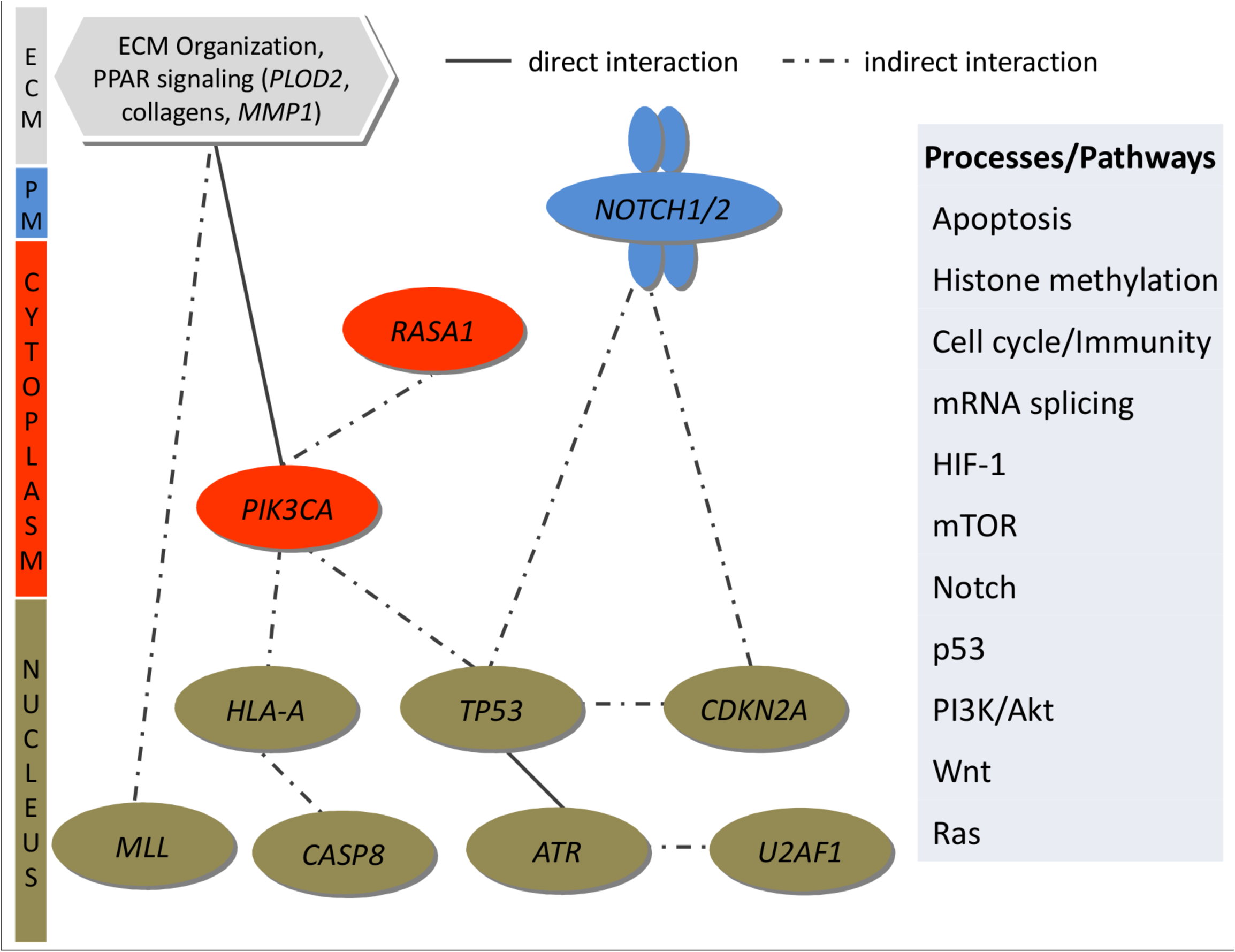
Significantly affected pathways in OTSCC. Genes harboring significant somatic variants and with expression changes in tumors were used in Cytoscape to derive a set of important signaling pathways implicated in OTSCC.

## Discussion

Squamous cell carcinomas of oral tongue are an aggressive group of tumors with a higher incidence in the younger population (≤50yrs), which spread early to lymph nodes and have a higher regional failure compared to gingivo-buccal cases [8-10]. Previous sequencing studies [3-5, 7] grouped oral tongue tumors with tumors from oral cavity but a rise in the incidence of oral tongue tumors, especially among younger people who never smoked, consumed alcohol or chewed tobacco, warrants further investigation of this subgroup of oral tumors. Additionally, the role of HPV in oral tongue tumors, unlike in oropharyngeal cases [18-20], is not well understood both in terms of incidence and prognosis. A meta-analysis of HPV-positive HNSCC tumors from multiple studies conducted at multiple locations concluded that HPV-positive patients, especially in oropharynx, have improved overall and disease-specific survival [21]. A past study has presented data that the HPV incidence in oral tongue is low [22] and some argue against any link between HPV infection and aggressive oral tongue tumors [23]. Although there is no consensus on rate of HPV incidence among oral tongue patients, it is generally believed that it is low compared to oropharyngeal tumors. However, some studies in the past [24], albeit from a different geography, established a much higher rate of HPV infection in oral tongue tumors.

We applied stringent filtering steps and used multiple annotation tools to come up with a list of 19 cancer-associated genes that harbored somatic variants in OTSCC. Most of these genes were also found in other studies, including the recent TCGA HNSCC study [7], with some notable differences. A comparison of somatic variants discovered in all HSNCC studies, including the current study, is provided in Additional file 12. The frequency for somatic changes in *CASP8, NOTCH1, CDKN2A* and *FAT1* genes in previous studies [3-7] were, 4-34%, 13-18%, 2-16% and 13-50%, respectively. This is different from what we found in the current study (8%, 4%, 6% and 2% respectively for the same genes). This may partly be attributed to the total number of tumor samples used in different studies but may also be due to a unique pattern of mutations specific to oral tongue subsite. It appears from our study that the latter is the case. For example, in one of the earlier studies [6] involving similar number of patients as in the current study, *CASP8* and *FAT1* were mutated in 34% and 50% of the patients but we find the frequency to be 8% and 2%, respectively. In some earlier studies, it was not possible to categorize and identify oral tongue-specific variants as the sites were classified under oral cavity [3, 4].

Although the somatic variants discovered from our study appear to be distributed uniformly across the genome, the significant copy number variation events are more concentrated in chromosomes 6-9 and 11 (Figure 1F and Additional file 8). One of the most important genes harboring somatic mutations discovered in our study is *CASP8*, the product for which derived from the precursor Procaspase-8. Caspase-8,is an important protein implicated in both apoptotic and non-apoptotic pathways [14]. Recent analysis from the TCGA study [7] suggests that mutations in *CASP8* co-occur with mutations in *HRAS,* and are mutually exclusive with amplifications in *FADD* gene. In our functional studies, the most important observation was that caspase-8 shows different effects in HPV-positive and HPV-negative cells, the effect being more pronounced in HPV-negative cells (Figure 4). Therefore, it is possible that HPV-negative tumors activate a completely different set(s) of pathways and/or may have different chemosensitivity towards drugs than the HPV-positive tumors. It was shown previously that HPV-positive HNSCC cell lines are resistant to TRAIL (tumor necrosis factor-related apoptosis-inducing ligand) and treatment of cells with the proteasome inhibitor bortezomib sensitizes HPV-positive cells towards TRAIL-induced cell death mediated by caspase-8 [25]. The E6 protein of HPV interacts with the DED domain of caspase-8 and induces its activation by recruiting it to the nucleus [26]. Our observation on the role of caspase-8-mediated apoptosis being more pronounced in the HPV-negative OTSCC cell line is similar to the observation on the role of *CASP8* in HPV-negative patients made earlier in TCGA study [7]. Taken together, genes including *CASP*8 regulate key pathways (Figure 6) that might play important role in the development of tumors in oral tongue.

In the past, several large sequencing studies have been undertaken in HNSCC [3-5, 7] that contained very few HPV-positive oral tongue patients. Our study is based on a unique patient cohort and attempts to link molecular signature with different clinical and epidemiological parameters. The prevalence of HPV is very high in oral tongue tumors from India, including in our cohort, compared with studies using cohorts elsewhere. We don’t see the same high prevalence of HPV in non-oral tongue tumors in oral cavity, for example in buccal tumors, in one of our study (Palve et al., *unpublished observation).* The exact reason for this high prevalence is not know. Additionally, the HPV positive patients that harbor mutations in *TP53* in some of the patient samples is also counter-intuitive owing to the fact that E6 is known to block p53. Although we don’t know the reason behind this, there is a possibility that HPV positive tumors harboring TP53 mutations represent a unique class of tumors and it will be interesting to see if those tumors recur early or late compared to the HPV positive tumors that have wild type p53 function. Therefore, this study is unique in that respect.

Identifying signature for tumor recurrence prospectively in primary tumors may add significant advantage to disease management. In order to do this, we used a machine learning method using the molecular changes identified in this study, in three batches of primary tumors; non-recurring, loco-regionally recurring and tumors with distant metastasis. We identified a 38-gene signature to be significantly distinguishing the three groups. The bootstrapping error for the non-recurring and the loco-regionally recurring groups were low (N = 34, *.632 error* = 0.03 and N = 10, *.632 error* = 0.3 respectively) but not in the metastatic tumor group (N = 4, *.632 error* = 1). This was due to the small sample numbers (N = 4) in the metastatic category, justifying the need for a larger sample set to validate the signature. The 38 gene signature identified in out study, however, needs to be validated in a much larger cohort in the future to achieve its true potential as a prognostic panel in OTSCC.

Finally, we were keen to see if the current study leads to finding novel drug candidates in OTSCC. We based our assumption on the fact that genome-wide somatic variant discovery in tumors may give rise to possibilities of finding novel drug targets/candidates or may led us to use existing drugs prescribed for other indications. In an attempt to identify if any of the significantly altered genes found in the current study could potentially act as drug targets, we screened for available drugs against them. We found drugs against three targets out of which two have undergone at least one clinical trial (Additional file 13).

## Methods

### Informed consent, ethics approval and patient samples used in the study

Informed consent was obtained voluntarily from each patient enrolled in the study. Ethics approval was obtained from the Institutional Ethics Committees of the Mazumdar Shaw Medical Centre. Matched control (blood and/or adjacent normal tissue) and tumor specimens were collected and used in the study. Only those patients, where the histological sections confirmed the presence of squamous cell carcinoma with at least 70% tumor cells in the specimen, were used in the current study. At the time of admission, patients were asked about the habits (chewing, smoking and/or alcohol consumption). Fifty treatment-naïve patients who underwent staging according to AJCC criteria, and curative intent treatment as per NCCN guideline involving surgery with or without post-operative adjuvant radiation or chemo-radiation at the Mazumdar Shaw Medical Centre were accrued for the study (Additional file 1). Post-treatment surveillance was carried out by clinical and radiographic examinations as per the NCCN guidelines.

### HPV detection

HPV was detected by using q-PCR using HPV16-and HPV18-specific TaqMan probes and primers and digital PCR using TaqMan probes and primers to detect HPV in primary tumor samples.

### Exome Sequencing, read QC, alignment, variant discovery and post-processing filters

Exome libraries were prepared using Agilent SureSelect, Illumina TruSeq and Nextera exome capture kits (Additional file 14) following manufacturers’ specifications. Paired end sequencing was performed using HiSeq 2500 or GAIIx and raw reads were generated using standard Illumina base caller. Read pairs were filtered using *in house* scripts (Additional files 15 and 16) and only those reads having ≥75% bases with ≥ 20 phred score and ≤ 15 Ns were used for sequence alignment against human hg19 reference genome using NovoAlign [27] (v3.00.05). The aligned files (*.sam) were processed using Samtools [28](v0.1.12a) and only uniquely mapped reads from NovoAlign were considered for variant calling. The alignments were pre-processed using GATK [29] (v1.2-62) in three steps before variant calling. First, the indels were realigned using the known indels from 1000G (phase1) data. Second, duplicates were removed using Picard (v1.39). Third, base quality recalibration was done using CountCovariates and TableRecalibration from GATK (v1.2-62), taking into account known SNPs and indels from dbSNP (build 138). Finally, UnifiedGenotyper from GATK (v2.5-2) was used for variant calling, using known SNPs and indels from dbSNP (build 138). Raw variants from GATK were filtered to only include the PASS variants (standard call confidence ≥ 50) within the merged exomic bait boundaries. Two out of 50 tumor samples did not confirm to the QC standards, therefore excluded from all further analyses. Therefore, all the downstream analyses were restricted to 48 primary tumors. The variants were further flagged as novel or present in either dbSNP138 or COSMIC (v67) databases, based on their overlap. In addition to GATK, we also used Dindel [30] to call indels. Both GATK and Dindel calls were filtered for microsatellite repeats (flagged as STR). The raw variant calls were used to estimate frequencies of nucleotide changes and transition:transversion (ti/tv) ratios. Exome-filtered PASS variants specific to the tumor samples, with respect to both location and actual call, were retained as somatic variants, which were further filtered to exclude variants where the region bearing the variant was not callable in the matched control sample, and those where the matched control sample had even one read covering the variant allele.

Scripts used to perform various filtering steps are provided in Additional file 16. The numbers of raw reads, after QC, alignment statistics, numbers of variants pre- and post-filters are provided in Additional file 2.

### Detection of cross-contamination and identification of significant somatic variants

We estimated cross-contamination using ContEst [31] in the tumor samples (Additional file 16). Locus-wise and gene-wise driver scores were estimated by CRAVAT [32] using the head and neck cancer database with the CHASM [33] analysis option. Genes with a CHASM score of at least 0.35 were considered significant for comparison with other functional analyses (Additional file 16). Somatic mutations were normalized with respect to the exome bait size (MB) to calculate the somatic mutation frequency per MB.

### Annotation and functional analyses of variants

Annotation and functional analyses of somatic variants was performed using IntoGen [34, 35](web version 2.4), MutSigCV [36, 37] and MuSiC2 [37]. Somatic variants, filtered to contain only those callable in the matched normal but not covered by any read in the control samples (VCF), were used for IntoGen with the ‘cohort analyses’ option. We also ran MutsigCV1.3 with these variants using coverage from un-filtered variants of all tumor samples (Additional file 16). Pooled alignments for all normal and tumor samples (bam), each, along with pooled variants for all normal samples (MAF) were analyzed using MuSiC2 to calculate the background mutation rates (bmrs) for all genes, and identify a list of significantly mutated genes (*p*-value of convolution test ≤ 0.05; Additional file 16). A condensed list of 19 genes, common between at least two analyses was compiled (Figure 1D).

### SNP genotyping and validation using Illumina whole-genome Omni LCG arrays

High quality DNA (200ng), quantified by Qubit (Invitrogen), was used as the starting material for whole-genome genotyping experiments following the manufacturer’s specifications. Briefly, the genomic DNA was denatured at room temperature (RT) for 10 mins using 0.1N NaOH, neutralized and used for whole genome amplification (WGA) under isothermal conditions, at 37^°^C for 20 hrs. Post WGA, the DNA was enzymatically fragmented at 37^°^C for 1hr. The fragmented DNA was precipitated with isopropanol at 4^°^C and resuspended in hybridization buffer. The samples were then denatured at 95^°^C for 20 mins, cooled at RT for 30 mins and 35*μ*l of each sample was loaded onto the Illumina HumanOmni 2.5-8 beadchip for hybridization (20hrs at 48^°^C) in a hybridization chamber. The unhybridized probes were washed away and the Chips (Human Omni2.5-8 v1.0 and v1.1, Additional file 2) were prepared for staining, single base extension and scanning using Illumina’s HiScan system.

We filtered the SNP locations to retain only those, called without any error, contained within the exome boundaries as per the sequencing baits, and which were callable (covered by at least five sequencing reads). At these locations, we estimated the overlap for individual SNP calls, i.e., chr/pos/ref/alt and for no calls; i.e., chr/pos/ref/ref; between sequencing and array platforms (Additional file 16).

### Discovering Copy number Variations (CNVs) and Loss of Heterozygosity (LOH)

CNVs and LOHs were identified using cnvPartition 3.1.6 plugin in Illumina GenomeStudio v2011.1, with default settings except for a minimum coverage of at least 10 probes per CNV/LOH with a confidence score threshhold of at least 100 (Additional file 17). Somatic CNVs and LOHs were extracted by filtering out any region common to CNVs and LOHs detected in its matched control. Somatic CNVs and LOHs were further filtered with respect to common and disease-related CNVs and LOHs using CNVAnnotator [38]. Overlaps with common CNVs and LOHs were discarded, reporting only the overlaps with disease-related, and novel CNVs and LOHs. We categorized the CNVs and LOHs within each cytoband and reported those with an occurrence in at least 10% of the patient samples.

### Gene Expression Assay

Gene expression profiling was carried out using Illumina HumanHT-12 v4 expression BeadChip (Illumina, San Diego, CA) in tumor and matched normal tissues (Additional file 9) following manufacturer’s specifications. Total RNA was extracted from 20mg of tissue using PureLink RNA (Invitrogen) and RNeasy (Qiagen) Mini kits. RNA quality was checked using Agilent Bioanalyzer using RNA Nano6000 chip. Samples with poor RIN numbers, indicating partial degradation of RNA, were processed using Illumina WGDASL assay as per manufacturer’s recommendations. The RNA samples with no degradation were labelled using Illumina TotalPrep RNA Amplification kit (Ambion) and processed according to the array manufacturer’s recommendations. Gene expression data was collected using Illumina’s HiScan and analyzed with the GenomeStudio (v2011.1 Gene Expression module 1.9.0) and all assay controls were checked to ensure quality of the assay and chip scanning. Raw signal intensities were exported from GenomeStudio for pre-processing and analyzed using R further.

Gene-wise expression intensities for tumor and matched control samples from GenomeStudio were transformed and normalized using VST (Variance Stabilizing Transformation) and LOESS methods, respectively, using the R package lumi [39]. The data was further batch-corrected using ComBat [40] (Additional file 16). The pre-processed intensities for tumor and matched control samples were subjected to differential expression analyses using the R package, limma [41] (Additional file 16). Genes with significant expression changes (adjusted *P* value < = 0.05) and fold change of at least 1.5 were followed up with further functional analyses.

### Recurrence prediction using random forests

We used presence or absence of somatic mutations/indels data in the entire set of genes for all the OTSCC patients, along with their recurrence patterns as training set for the random forests [11] analyses using the varSelRF package in R. This method performs both backward elimination of variables and selection based on their importance spectrum, and predicts recurrence patterns in the same set by iteratively eliminating 2% of the least important predictive variables until the current OOB (out-of-bag) error rate becomes larger than the initial or previous OOB error rates. In order to understand the specificity of the best minimalistic predictors of tumor recurrence, we estimated the 0.632+ error rate [17] over 50 bootstrap replicates. We used the varSelRFBoot function from the varSelRF Bioconductor package to perform bootstrapping. The .632+ method is described by the following formula:

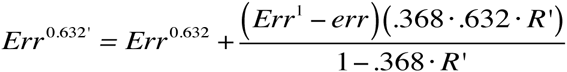

where *Err*^(.632’)^, *Err*^(.632)^*, Err*^(1)^ and *err* are errors estimated by the .632+ method, the original .632 method, *leave-one-out* bootstrap method and *err* represents the error. *R’* represents a value between 0 and 1. Another popular error correction method used is *leave-one-out* bootstrap method. The .632+ method was designed to correct the upward bias in the *leave-one-out* and the downward bias in the original .632 bootstrap methods.

For all iterations of all random forest analyses, we confirmed that the variable importance remained the same before and after correcting for multiple hypotheses comparisons using pre- and post-Benjamin-Hochberg FDR-corrected *P* values (data not shown). R commands for variable elimination using random forests, 0.632+ bootstrapping and re-computing importance values after multiple comparisons testing are provided in Additional file 16.

### Pathway analyses

Consensus list of genes from analysis, filtering and annotation of variant calls and from differential expression analysis using whole genome micro-arrays, were mapped to pathways using the web version of Graphite Web [42] employing KEGG and Reactome databases. The network of interactions between genes was drawn originally using CytoScape [43] (v3.1.1) using the .sif file created by Graphite Web (Additional file 16).

### Data Visualization

We used Circos [44] (v0.66) (Additional files 18 and 19) for multi-dimensional data visualization. Additionally, we used the cbioportal protal (http://www.cbioportal.org/) to visualize variants within the 19 genes harboring significant variants. All of the mandatory fields accepted by Mutation Mapper were provided for select genes from our study to create structural representations for each gene including domains. Such diagrams from our study, the HNSCC study and all cancer studies from TCGA were collated using the image-editing tool, GIMP (www.gimp.org). SNPs and indels were visualized for each individual tumor sample using IGV [45] (v1.5.54), along with the reads supporting variants (Additional file 20).

### Validation of somatic variants using Sanger sequencing

Primers were designed using the NCBI primer designing tool (http://www.ncbi.nlm.nih.gov/tools/primer-blast/index.cgi?LINK_LOC=BlastHome) and used in Sanger sequencing for validation. The sequences of all primers (IDT) used for validation is provided in Additional file 21. We tested the specificity of the designed primers using UCSC’s tool, *In Silico* PCR. The variant-bearing region was amplified by using specific primers and used in Sanger sequencing (Additional file 14). The somatic variants were confirmed by sequencing in the entire tumor and matched control DNA set used for the exome sequencing followed by further validation in 60 additional tumor samples (Additional file 1B).

### Cell culture and knockdown of CASP8 gene

The human OTSCC cell lines UPCI:SCC040 (gift from Dr. Susan Gollin, University of Pittsburgh, PA, USA) [16] and UM-SCC47 (gift from Dr. Thomas Carey, University of Michigan, MI, USA) [15] were used in the study. All the cells were maintained in Dulbecco’s Modified Eagles’ Media (DMEM) supplemented with 10% FBS, 1% MEM nonessential amino acids solution & 1% penicillin/streptomycin mixture (Gibco) at 37°C with 5% CO2 incubator.

We performed the siRNA-based knockdown using UPCI:SCC040 and UM:SCC47 cell lines for *CASP8* gene. The expression of Caspase-8 was transiently knocked down using ON-TARGETplus Human *CASP8* smart pool siRNA (L-003466-00-0010; Dharmacon) along with an ON-TARGET plus Non-targeting siRNA (D-001810-01-20; Dharmacon). The transfection efficiency for the two cell lines (UPCI:SCC040 and UM:SCC47) were optimized using siGLO Red Transfection Indicator (D-001630; Dharmacon). The siRNA duplexes were transfected using Lipofectamine-2000 according to the manufacturer’s instructions (Invitrogen). The siRNA-oligo complexes medium was changed 8 hrs post transfection. The efficiency of transfection along with the mRNA expression was analyzed at 24 and 48 hrs post transfection by qRT-PCR. The specific down-regulation of *CASP8* was confirmed by three independent experiments.

### RNA isolation and quantitative real-time PCR

RNA was extracted from cell pellets and tissues using RNeasy Mini kit spin columns (Qiagen) following manufacturer’s protocol. Genomic DNA contamination was removed by RNase-Free DNase Set (Qiagen) and the total RNA was eluted in nuclease free water (Ambion). The RNA samples were estimated using Qubit 2.0 fluorometer (Invitrogen) and the integrity was checked by gel electrophoresis. The RNA samples were stored at -80^°^C until further used. The cDNA was synthesized with 400ng total RNA, using a SuperScript-III first strand cDNA synthesis kit, and following the manufacturer’s instructions (Invitrogen). The cDNA was then subjected for quantitative real-time PCR (q-RT-PCR) using KAPA SYBR FAST qPCR Master Mix (KK4601, KAPA). The primer pairs used for testing the expression of caspase-8 in q-RT-PCR were, forward 5’-ATGATGACATGAACCTGCTGGA-3’ and reverse 5’-CAGGCTCTTGTTGATTTGGGC-3’. The amplification was done on Stratagene MX300P real time machine. To normalize inter-sample variation in RNA input, the expression values were normalized with GAPDH. All amplification reactions were done in triplicates, using nuclease free water as negative controls. The differential gene expression was calculated by using the comparative C_T_ method of relative quantification [46].

### Assessment of cell viability (confirm)

MTT cell proliferation assay was performed as per manufacturer’s instructions (Sigma) to assess cell viability. Briefly, cells were seeded on 96-well plates containing DMEM with 10% FBS & incubated overnight. After treatment with 0.1% DMSO (vehicle control), or Cisplatin for 48 hrs, medium was changed and 100 *μ*l of MTT solution (1mg/ml) was added to each well. The cells were further incubated for 4hrs at 37°C. The formazan crystals in viable cells were dissolved by adding 100*μ*l of dimethyl sulfoxide (DMSO) (Merck). The absorbance was recorded at 540 nm using reference wavelength of 690 nm on micro plate reader (Tecan Systems). Data were normalized to vehicle treatment, and the cell viability was calculated using GraphPad Prism software (version 4.03; La Jolla, CA). All the experiments were performed in triplicates.

### Wound healing assay

Cells were cultured up to 80% confluency in 12 well plates; serum-starved for 24 hrs and then wounded using a 200*μ*l pipette tip. The wound was washed with 1x PBS and the cells were grown in DMEM containing 10% FBS. Cells were imaged at 10x magnification at 0 hr, 15 hrs, 23 hrs and 42 hrs. For each well, three wounds were made and the migration distance was photographed and measured using Carl Zeiss software (Zeiss). Each experiment was performed in triplicates.

### Matrigel invasion assay

The ECM gel (E1270, Sigma) was thawed overnight at 4^o^C and plated at requisite concentrations (for UPCI:SCC040: 1.5mg/ml and UMSCC047: 2mg/ml) onto the transwell inserts and incubated overnight in the CO_2_ incubator at 37^o^C with 5% CO_2_. Cells were serum-starved for overnight, harvested, counted and seeded (UPCI:SCC040: 50,000 cells and UMSCC047: 20,000 cells per well) on top of the matrigel transwell-inserts (2 mg/ml) in serum free medium as per manufacturer’s specifications (Sigma). D-MEM containing 10% FBS and 1% NEAA was added to the lower chamber. The 24-well plates containing matrigel inserts with cells were incubated in 37°C incubator for 48 hrs. At the end of incubation time, cells in the upper chamber were removed with cotton swabs and cells that invaded the Matrigel to the lower surface of the insert were fixed with 4% paraformaldehyde (Merk Milipore), permeabilized with 100% methanol, stained with Giemsa (Sigma), mounted on glass slides with DPX mounting agent and counted under a light microscope (Zeiss). Each experiment was performed in triplicates.

## Conclusions

We have catalogued genetic variants (somatic mutations, indels, CNVs and LOHs) and transcriptomic (significantly up- and down-regulated genes) changes in oral tongue squamous cell carcinoma (OTSCC) and used those in an integrated approach linking genes harboring somatic variants with common risk factors like tobacco and alcohol; clinical, epidemiological factors like tumor grade and HPV; and tumor recurrence. We found *CASP8* gene to be significantly altered and play an important role in apoptosis-mediated cell death in an HPV-negative OTSCC cell line. Finally, we present data towards a minimal gene signature that can predict tumor recurrence.

## Availability of supporting data

All supporting data is available through the journal’s website and on figshare.

**List of abbreviations:** Oral tongue squamous cell carcinoma (OTSCC), head and neck squamous cell carcinoma (HNSCC), single nucleotide variant (SNV), insertion and deletion (indel), loss of heterozygosity (LOH), copy number variation (CNV)

## Competing interests

None

## Authors’ contributions

BP: conceived, designed and supervised the study, wrote the manuscript; NMK: analyzed the data and wrote the manuscript; SG: analyzed the data and critically read the manuscript; SP, PJ, CK, VKP: analyzed the data; VP, GS, AS: produced data on *CASP8* functional analysis; LV, AKH, MP: produced sequencing data; KD and JN: produced array data; GS, AS, VK and MAK: provided clinical data, clinical input and associated clinical information.

## Acknowledgements

Research presented in this article is funded by Department of Electronics and Information Technology, Government of India (Ref No: 18(4)/2010-E-Infra., 31-03-2010) and Department of IT, BT and ST, Government of Karnataka, India (Ref No:3451-00-090-2-22).

## Additional files

All the additional files can be downloaded from figshare

(http://figshare.com/s/c928faa66f2a11e586d506ec4b8d1f61).

Additional file 1:

Title: Patient details used in the study.

Additional file 2:

Title: Sequencing, read QC, alignment and variant calls, OMNI SNP genotyping array validation.

Additional file 3:

Title: Validation using capillary gel electrophoresis based on Sanger sequencing in A. discovery set and B. validation set.

Additional file 4:

Title: The ratio of transitions to transversions (ti/tv) was estimated using the exome-filtered GATK PASS variants for tumor and matched control samples. The dotted lines depict the respective median ti/tv ratios.

Additional file 5:

Title: Effect of habits, clinical parameters and HPV infection on individual nucleotide change.

Additional file 6:

Title: Functional annotation of somatic variants using IntOGen, MuSiC2, MutSigCV.

Additional file 7:

Title: Position and frequency of somatic variants in protein domains found in this study, TCGA HNSCC, and in studies involving all cancer types using mutation mapper in the cBioPortal.

Additional file 8:

Title: Cytoband-wise representation of CNVs found in all 48 samples along with clinical parameters and patient epidemiology.

Additional file 9:

Title: Transformed, normalized and batch-corrected intensities following expression assay and results from differential expression analyses.

Additional file 10:

Title: Functional validation for the role of *CASP8* in OTSCC cell lines.

Additional file 11:

Title: List of all pathways affected by somatic mutations/indels, copy number variations and expression changes (log_2_FC≥=0.6).

Additional file 12:

Title: Comparative sample frequency of important variants found in this and other HNSCC studies.

Additional file 13:

Title: Drug candidates and their targets in head and neck cancer.

Additional file 14:

Title: Supplementary Methods.

Additional file 15:

Title: Read QC filter scripts’ executable.

Additional file 16:

Title: Scripts used in the study.

Additional file 17:

Title: GenomeStudio output of all LOHs and CNVs found using cnvPartition plugin in GenomeStudio.

Additional file 18:

Title: Circos data and config files.

Additional file 19:

Title: Circular genomic representation using Circos (v0.66) of LOHs, somatic variants, CNVs with > = 10% frequency of patients bearing them, and genes with significant expression changes (|log_2_FC|>=0.6).

Additional file 20:

Title: IGV snapshots of all significantly mutated somatic variants in this study.

Additional file 21:

Title: Primer sequences used in the Sanger validation study.

